# Genome-wide association study reveals putative bacterial risk factors for cavitary *Mycobacterium avium* complex lung disease

**DOI:** 10.1101/2021.07.06.451401

**Authors:** Hirokazu Yano, Yukiko Nishiuchi, Kentaro Arikawa, Atsushi Ota, Mari Miki, Fumito Maruyama, Hiroshi Kida, Seigo Kitada, Tomotada Iwamoto

## Abstract

*Mycobacterium avium* complex (MAC) lung disease is a slowly progressive disease, and its increasing incidence has garnered increased research interests. Cavitary MAC lung disease is associated with a higher mortality rate. Though genetic studies have unraveled the human risk factors, the role of microbial factors on pathogenesis behind the disease remains elusive. In this study, *M. avium* isolates were collected from sputum specimens of 109 distinct Japanese patients with or without a cavity (60 with a cavity and 49 without cavity) in a hospital located in Osaka prefecture. *M. avium* genomes were sequenced and searched for DNA motifs associated with cavity formation using a bacterial GWAS. Excluding known macrolide resistance mutations; cavity formation was found to be primarily associated with variants of cytochrome P450 of the CYP139 family, type I polyketide synthase Pks13, and the promoter region of an operon encoding membrane-anchored protease FtsH and folate synthesis pathway enzymes. Cavity risk variants at these three loci were frequent in the MahEastAsia2 lineage among the six lineages detected in *M. avium* global populations. Furthermore, the study demonstrated a correlation between the cavity risk promoter variant and increased sulfamethoxazole**/**trimethoprim resistance. Together, these findings suggest that natural variation in the biosynthesis and maintenance processes of *M. avium* membrane components influences the disease type of MAC lung disease. Although further validation is needed, the bacterial genetic markers listed in the present study could contribute to prognosis prediction based on bacterial genotyping and help develop treatment strategies in the future.

**IMPORTANCE:** Nontuberculous mycobacterial lung disease is of great concern in countries with an increasingly aging population. The disease types can largely be classified into non-cavitary nodular bronchiectasis and cavitary diseases (fibrocavitary, nodular bronchiectasis with cavity) that require different treatment strategies depending on the causal agents. Several studies have reported human risk factors for the disease; however, little efforts were made to investigate the risk factors in nontuberculous mycobacteria. Moreover, molecular genetics experiments have been difficult to search for virulence factors in *M. avium*, which the population genomics approaches could overcome. Here, the GWAS results suggested variants in three chromosomal loci associated with mycobacterial membrane components as risk factors for cavitary MAC lung disease. These findings could help develop treatment strategies for MAC lung disease in the future.

## INTRODUCTION

Nontuberculous mycobacterial (NTM) lung disease is a slowly progressive disease caused mainly by *Mycobacterium avium* complex (MAC) that are ubiquitously present in the environment (1–5). The increasing incidence of the disease has become a global threat; moreover, the disease varies with ethnicity, showing a higher incidence in Asian countries than in Caucasian countries (6–9). Studies have reported that several human risk factors, particularly the underlying disorders predispose to NTM lung disease (1, 10). In addition, a recent genome-wide association study (GWAS) has revealed that NTM lung disease is associated with expression of calcineurin like EF-hand protein 2 (CHP2) encoded on chromosome 16p21 (11). These reports suggest that human genetic factors, at least in part, are involved in the onset of the disease; however, the precise role of bacterial genetic factors on the onset and severity of the disease remains elusive.

The MAC consists of *M. avium, M. intracellualre, M. chimaera*, and nine other less frequently reported taxa (12). Among MAC members, *M. avium* is the most frequently detected species in clinical samples of NTM lung disease in Japan and the USA (13, 14). Although *M. avium* was further classified into multiple subspecies, subsp. *hominissuis* (MAH) is the only subspecies identified in human clinical samples (15). Comparative genomics analyses between global MAH strains have reported considerable genetic distances between East Asian isolates and the rest (16–20). For example, we previously identified five major lineages, of which two lineages, MahEastAsia1 (EA1) and MahEastAsia2 (EA2), were predominant in East Asian isolates while elucidating the genetic population structure of global MAH population (19). In addition, studies have also revealed that the MAH genomes possess many recombination footprints, some of which are unique to the strains, suggesting that chromosome recombination among MAH lineages is not just an ancient event but is ongoing in nature unlike *M. tuberculosis* or *M. avium* subsp. *paratuberculosis* (19, 21, 22). As genome-wide recombination shuffle alleles in local population, we could speculate that there might be a considerable phenotype variation in local MAH population. The phenotype variation may be seen in virulence levels, and thereby affect disease type of patients.

MAC lung disease can clinically be classified into non-cavitary nodular bronchiectatic disease (bronchiectasis) (NB) and cavitary disease. NB, a slowly progressive disease (23), is characterized by bronchial dilatations with multiple small nodules (1). Cavitary disease, represented by the fibrocavitary (FC) disease, but not fully classified, is a rather progressive disease of the cavity. The FC disease is characterized by the cavity accompanying fibrosis, mostly found in the upper lobes (1). FC predominantly occurs in people with a history of smoking, while NB occurs mostly in nonsmoking postmenopausal females (24); however, NB can coexist with the cavity in the lung parenchyma or with FC (25, 26). It has been shown that the cavity volume is negatively correlated with lung function (26), and the patients diagnosed with cavities (FC or NB with cavity) had a higher mortality rate than those without (25.1% versus 0.8%) (27). Therefore, more aggressive treatment approaches are recommended for cavitary disease than NB (28). Hence, understanding the risk factors in the bacterial genome for cavitary lung disease is a prerequisite for the effective management of MAC lung diseases.

Genetic studies based on exome sequencing or GWAS have reported causal loci for the host susceptibility of MAC lung diseases (11, 29). However, the specific bacterial factors associated with the onset of MAC lung disease or the induction of cavitary MAC lung diseases have not been explored, which could partly be due to the lack of a bacterial strain set linked to the host’s information, and in part due to the lack of appropriate animal infection models for MAC lung disease. Recently developed molecular genetics (TnSeq) systems applied to two models, *M. avium* strains, MAC 109 and 11, has provided mechanistic insights into the virulence and antibiotic resistance of MAH (30, 31). While the TnSeq approach contributes to understanding the fundamental process of bacterial fitness under various controlled conditions, it has limitations in understanding variations in natural populations at the original bacterial niche. Therefore, we aimed to complement the MAC lung disease research by adding information on the bacteriological aspect using a population genomics approach.

Bacterial GWAS have revolutionized understanding the genetic bases of virulence or antimicrobial resistance in pathogens (32–36). However, GWAS to detect the genetic variants associated with pathogenicity of NTM in lung disease has not been explored. Therefore, in this study, we build a clinical *M. avium* dataset to link the bacterial genome with disease type, year of isolation, and history of antimicrobial therapy. Further, this study investigates the bacterial causal loci for cavitary MAC lung disease employing GWAS.

## RESULTS

### *M. avium* Osaka clinical dataset

*M. avium* strains used in GWAS were collected from 109 distinct Japanese adults and immunocompetent individuals diagnosed with a pulmonary disease commuting to National Hospital Organization Osaka Toneyama Medical Center (Toyonaka, Osaka, Japan). Among 109 patients, 49 patients possessed NB alone or unclassifiable disease type without a cavity, while 60 patients possessed cavity (Fig. 1, Dataset S1). Fifty-seven patients with cavities and 17 without cavities had a history of antimicrobial treatment before the bacterial strain isolation.

**FIG 1.**
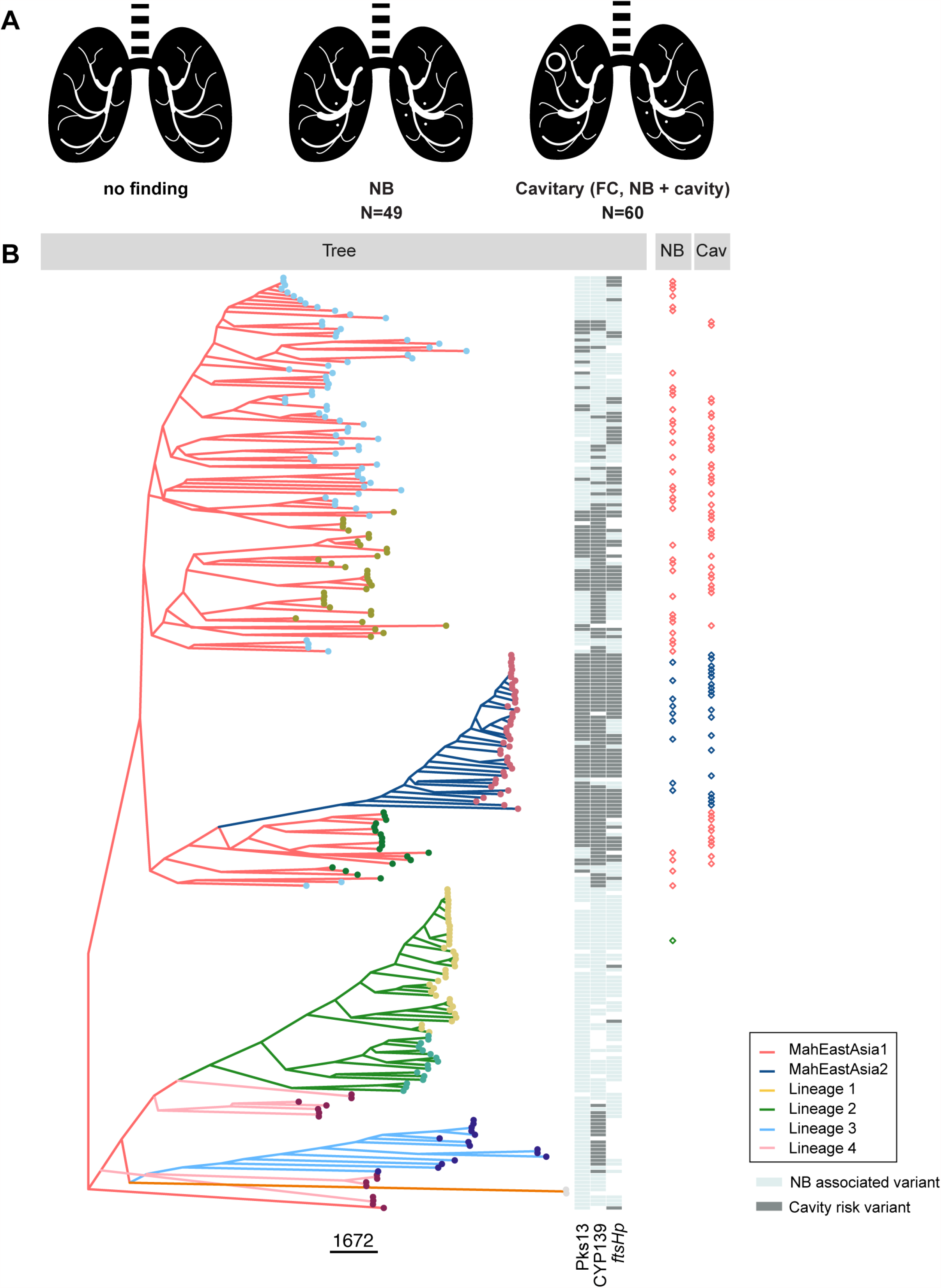
*M. avium* Osaka strains. **(A)** Typical characteristics of MAC lung diseases: the non-cavitary disease (mainly NB) and cavitary diseases. (**B)** Neighbor-joining tree of 255 global strains. The distance used is p-distance. Tip color indicates sequence cluster (SC), while the branch color indicates the lineage to which each strain belongs. The disease type linked to 109 Osaka strains and the variant type of three focused loci are indicated next to the tips. The variants searched for are as follows: Pks13, unitig n194039 for NB, n194036 for cavity; CYP139, D201-N220 for NB, N201-S220 for cavity; *ftsHp*, n218639 (+) n193112 (-) for NB, n218639 (-) n193112 (+) for cavity. Strains carrying neither NB nor cavity-associated variant were left blank in the heatmap. Tree with tip label is provided in figshare DOI:10.6084/m9.figshare.14788923.

As *M. avium* strains are all from Japanese patients, it was expected that most strains belonged to either of the two East Asian lineages (19). The lineage of these strains was inferred using a genomic dataset of 255 strains containing global isolates based on fastGEAR analysis (37), in which the lineage has been defined by hierarchical clustering based on the proportion of genomic segments sharing the ancestry between two genomes using linkage information (Fig. S1, Dataset S2). Six lineages— EA1, EA2, lineage 1, lineage 2, lineage 3, and lineage 4—were detected in the 255 global strains (Fig. 1B). As expected, most Osaka strains fell into EA1 or EA2 lineages. One strain, PPE_S61, fell into lineage 2 (previously called SC2), where strain 104 (a widely used model strain) and most European strains belong. EA1 and EA2, mostly formed by Japanese clinical strains, also included several USA strains (Dataset S2). Lineage 4 (previously called SC4) genomes strikingly carried large chromosomal chunks from lineage 2 and EA1 (Fig. S1).

### Overview of GWAS results

Since the core genomic region covered only 76% of each draft genome, even for the Osaka dataset, we chose to find an association of unitigs (graph node in compacted De Bruijn Graph of k-mers) instead of core genomic SNPs. Furthermore, EA2 genomes have been shown to be markedly different from EA1 genomes, do not possess many recombination footprints (21); therefore, we used a linear mixed model method implemented in the DBGWAS pipeline (38) to control for population stratification in the bacterial dataset.

A total of 47,503 unique presence/absence patterns of unitigs with frequencies > 5% in the population were used for association analysis in the linear mixed model. The Manhattan plot and Q-Q plot showing the *P* value for locus effect (referred to as weight in DBGWAS output) are shown in Fig. 2A and 2B, respectively, where the Q-Q plot indicates the absence of *P* value inflation. Based on the Q-Q plot results, unitigs with a *P* value < 10^−4^ were identified as candidates for NB (no cavity) or cavity-associated variants, wherein the Manhattan plot identified five major *P* value peaks (Fig. 2A). These unitigs were found to be located in the 23S rRNA gene, the promoter region of *ftsH*, and the protein coding regions of dihydrodipicolinate reductase, CYP139 family cytochrome P450, and type I polyketide synthase Pks13. The lineage effect was not significant for the principal components detected in the analysis.

**FIG 2.**
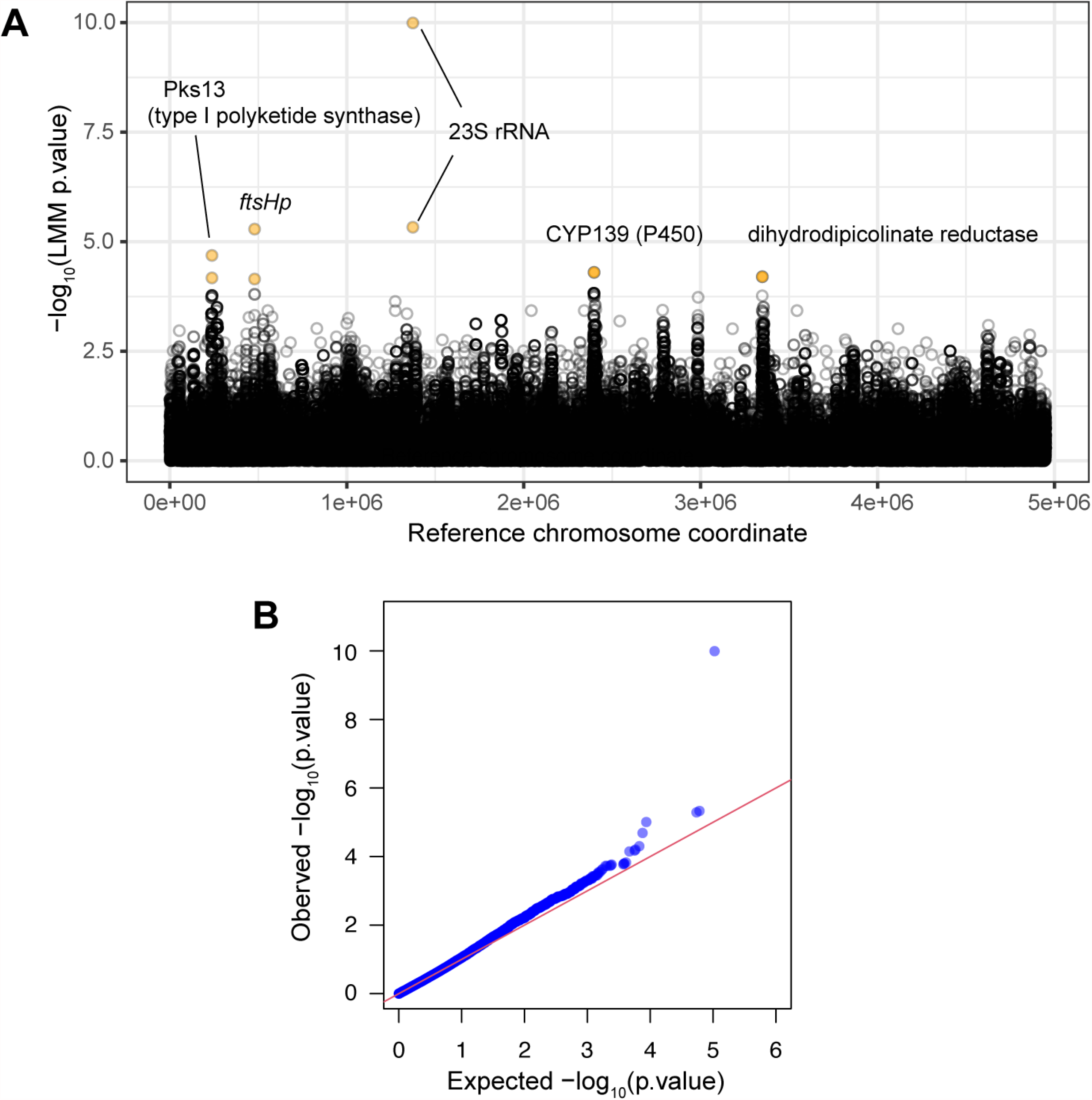
Overview of GWAS results. **(A)** Manhattan plot. The plot shows the *P* value assigned to each unitig matched to strain TH135 chromosome. **(B)** Q-Q plot. *P* values for 47,503 unique unitig patterns are shown. X-axis values are generated by random sampling from a uniform distribution.

### 23S rRNA gene

Unitigs in the 23S rRNA gene (*rrl*) included the two major alleles (*P* = 1.0e-10, *P* = 4.6e-06) with negative effect size value, three minor alleles with positive effect size value (*P* = 1.4e-03 to 6.1e-01), and two rare alleles (allele frequency; af < 6) were filtered out before regression analysis (Comp_0, in Dataset S3). All five minor alleles were known clarithromycin resistance mutations: A2058G, A2058C, A2058T, A2059G, and A2059C (numbers are equivalent positions in *E. coli rrl*) (39, 40). These are unlikely to be causal mutations in cavity formation and, therefore, are out of focus in this study.

### Dihydrodipicolinate reductase

Although clusters of unitigs with low *P* values were obvious in the coding region of dihydrodipicolinate reductase (locus_tag MAH_3077; coordinate 3,348,629), nonsynonymous substitution was not detected in the biallelic unitig pair with the lowest *P* value (unitig ID: n34259 and n74152, *P* = 6.29e-5) in this region. Moreover, the synonymous substitution around this coordinate affecting the bacterial phenotype through gene expression or translation is unknown. Therefore, we speculated that the significant locus effect detected for these unitigs could be due to the hitchhiking effect. However, the causal mutation, which may be located in neighboring regions, could not be identified in this study. Therefore, we focused on the features of the remaining three *P* value peaks.

### *FtsH* promoter region

Two significant unitigs matching the reference chromosome coordinates 478,982 to 479,033 are found in the intergenic region between MAH_0464 locus encoding the VOC family protein and MAH_0465 locus (*ftsH*) encoding ATP-dependent zinc metalloprotease. The *fol* genes that encode folate synthesis pathway enzymes are the immediate downstream of *ftsH* (Fig. 3A). All genomes analyzed carried a direct repeat of a 39 bp sequence in this region (hereafter referred as to the *ftsH* promoter region); however, the repeat number and the repeat sequences varied among the genomes (Fig. 3B, Fig. S2 for alignment). One repeat variant carried T at the 9th position of the 39 bp sequence, while the other variant carried C at the equivalent position (Fig. 3B and 3C). Five unitigs with *P* < 10^−3^ overlapping with this region are shown in Fig. 3D. All of these contained either the full-length or a part of the two 39 bp repeat variants. The unitigs carrying the C variant showed a positive effect size value, indicating an association with the cavity.

**FIG 3.**
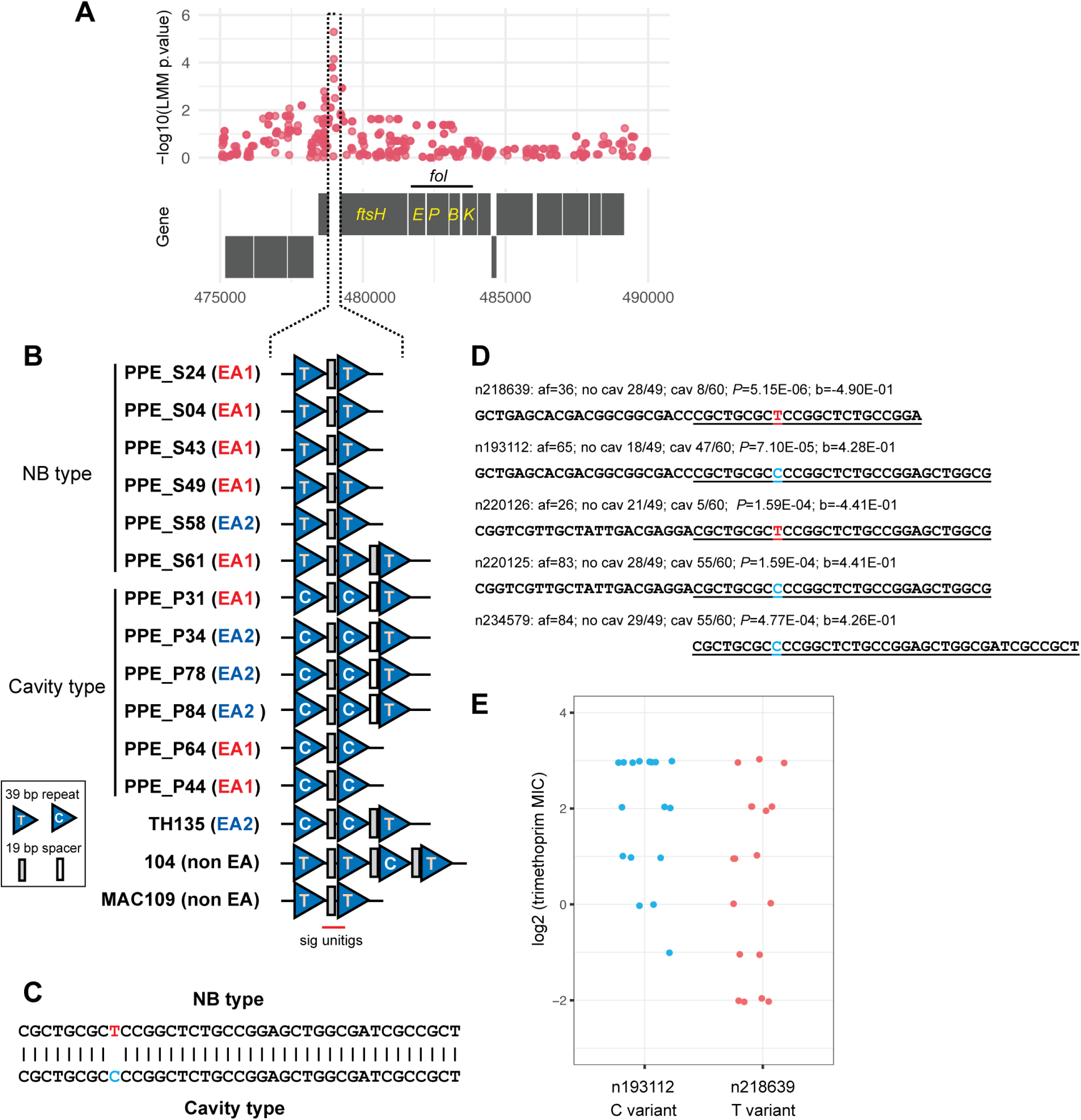
Association of the *ftsH* promoter variants with cell phenotype and host disease. **(A)** Locus effect *P* values around the *ftsH* promoter region. (**B**) Schematic figure showing repeat organization in the *ftsH* promoter region. Note that the 19 bp spacer sequence is present between the 39 bp repeat. The alignment is shown in Fig. S2. (**C**) Nucleotide sequences of two repeat unit variants. (**D**) Significant unitigs are overlapping with the 39 bp repeat. Unitig information is as follows: af, allele frequency; *P, P* value; b, effect size (weight in DBGWAS); NB, frequency in no cavity group; cav, frequency in cavity group. Sequence overlapping with either of the two 39 bp repeat variants is underlined. n218639 and n193112 spans the first and the second repeats from the left on panel B. Note that strain 104 genome carries both n218639 and n193112 as shown in panel B. (**E**) MIC of sulfamethoxazole/trimethoprim mix of Osaka strains. Y-axis value is log_2_ transformed value of trimethoprim concentration in the mix. N = 17 for the C variant, n = 18 for the T variant. MIC = 8 denotes that MIC is equal to or greater than 8. The difference between the two groups is significant in the non-parametric Wilcoxon-Mann-Whitney test (*P* = 0.02564)

To confirm the association of this promoter with the cell phenotype, we investigated the susceptibility of Osaka strains to sulfamethoxazole/trimethoprim mix that target folate synthesis. The rationale is that if the *ftsH* promoter is more active in one group, the Fol enzymes should be produced more in the group, and therefore, one group should show higher resistance to the sulfamethoxazole/trimethoprim mix. Nineteen strains carrying the T variant (unitig ID n218639 in Fig. 3D) and 16 strains carrying the C variant (n193112 in Fig. 3D) were selected, and their respective minimum inhibitory concentrations (MICs) were measured. A few strains carrying both n218639 and n193112 (e.g., strain 104 in Fig. 3B) were present in the Osaka strain set, and this genotype was more frequent in the global population (Dataset S2). These were excluded from the experimental analysis. Although the variation within the group was large, the mean MIC was significantly higher in the C variant group than in the T variant group (Z = 2.2144, *P* = 0.02564, in Exact Wilcoxon-Mann-Whitney Test) (Fig. 3E). These results revealed that the folate synthesis pathway of the C variant group is less susceptible to sulfamethoxazole/trimethoprim mix than the T variant group and indicated that the C variant could be associated with a higher promoter activity, leading to increased expression of *ftsH* and *fol* genes, than the T variant.

### Cytochrome P450 of the CYP139 family

A cluster of unitigs with low *P* values was located in the coding region of one specific cytochrome P450 homologue (locus_tag MAH_2197 in TH135, MAV_3106 in 104; DFS55_11100 in MAC 109) (Fig. 4A), belonging to the CYP139 family with an undetermined biological role (41). The lowest *P* value in this region was found on two co-occurring unitigs: one (n98104) from TH135 coordinates 2,397,713 to 2,397,773 coding for 13 amino acids, and the other (n234809) from 2,397,793 to 2,397,831 coding for 20 amino acids (Fig. 4A, comp_33 in Dataset S3). Both have the same af of 34 and are associated with no cavity (*P* = 5.0e-5, effect size = -4.9e-1) (Fig. 4B). Additionally, two unitigs with positive effect size, coding for an alternate amino acid at position 201 or 220 of the gene product, were also identified. Two alternate amino acids at positions 201 and 220 were found in three patterns in the *M. avium* population: N201 (codon AAC) -S220 (AGC), D201 (GAC) - S220 (AGC), and D201 (GAC) -N220 (AAC) (Dataset S2); however, a combination of D201-S220 was rare (2/109) in the Osaka strains. The frequencies of D201-N220 and N201-S220 showed a different distribution between the no cavity and the cavity groups (D201-N220: 26/48 in no cavity, 8/59 in cavity; N201-S220: 22/48 in no cavity, 51/59 in cavity) in a subset of Osaka strains without D201-S220 (Chi^2^ = 18.302, df = 1, *P* = 1.88e-05). These results suggested that the specific amino acid positions of CYP139 could be associated with disease type. Furthermore, amino acids at positions 201 and 220 constitute a part of the loop located on the surface of the same side of the three-dimensional structure of CYP120A1 family P450 (Fig. 4C), implying their cooperative roles in the mechanisms of action.

**FIG 4.**
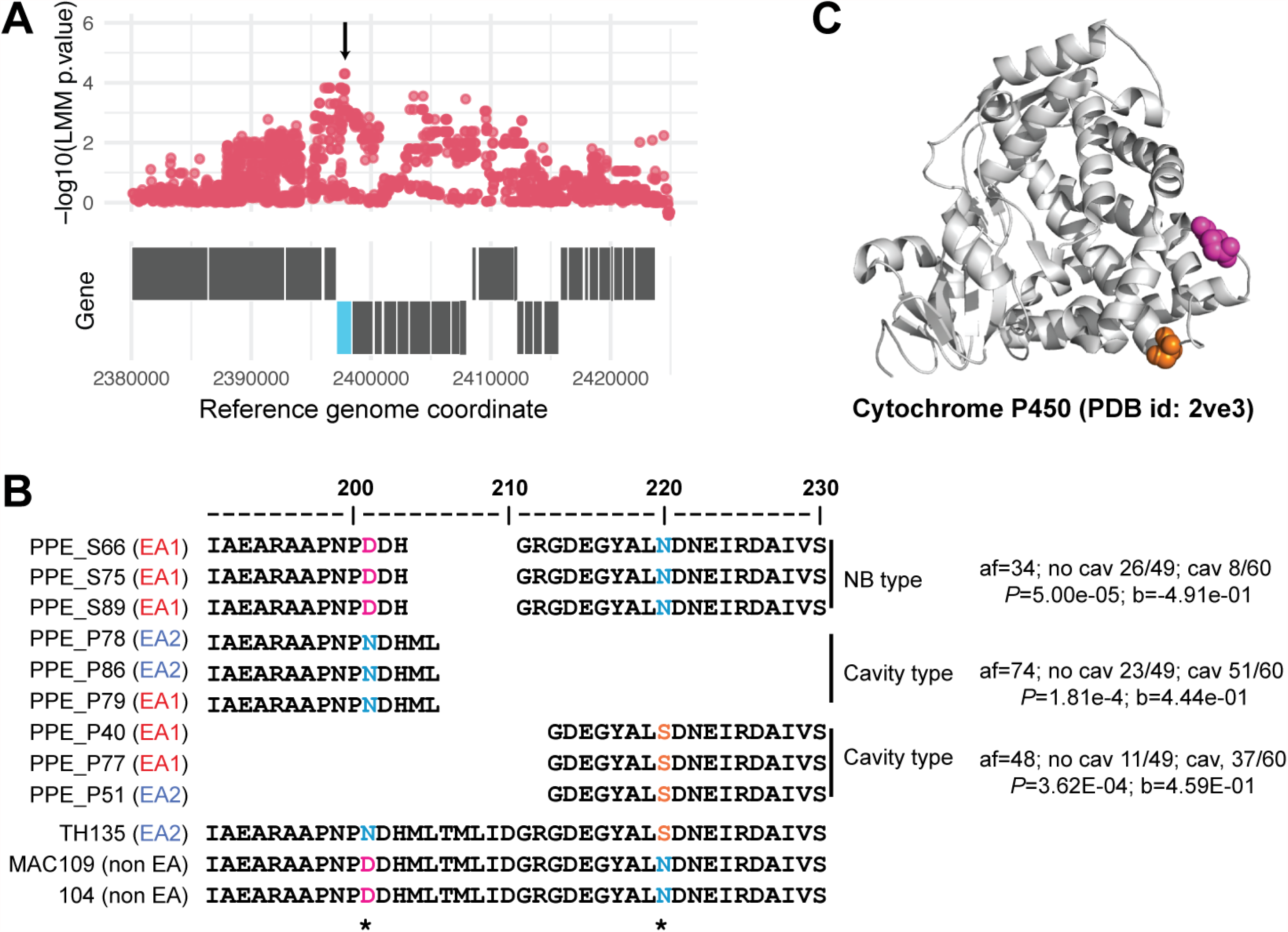
The disease type-associated amino acids in CYP139. (**A**) Locus effect *P* values around coordinate 2,400,000. (**B**) Amino acid sequences coded on significant unitigs in representative genomes. (**C**) Positions of N201 and S220 equivalent amino acids in the three-dimensional structure of cytochrome P450 belong to the CYP120A1 family (PDB id 2ve3) (73). N201 and S220 equivalent positions on the PDB database proteins were identified by generating alignment in JPred 4 (74) using the product of MAH_2197 as a query. N201 and S220 equivalent amino acids (E221, S240) on CYP120A1 were highlighted using PyMol v.2.4.0 (Schrödinger, Inc., New York, USA). Magenta, N201; Orange, S220.

### Type I polyketide synthase Pks13

The *P* value peak around the coordinate 239,000 shown in Fig. 2A was identified to be within the coding sequence of a type I polyketide synthase (locus_tag MAH_0232 in TH135, MAV_0218 in 104, DFS55_23815 in MAC 109) (Fig. 5A), an ortholog of Pks13 (Rv3800c) of *M. tuberculosis* (42), which has attracted attention as a target of new antitubercular drugs (43–45). Psk13 plays pivotal roles in the condensation of mycolic acid assembly (42, 46) and trehalose monomycolate (TMM) synthesis (47). The top-ranked unitig (n194039, *P* = 2.05E-05) was associated with a negative effect size value (thus no cavity), and its opposed unitig (n194036) contained only one synonymous substitution (Fig. 5B). The 15th ranked unitig (n126875, *P* = 4.94E-04, Dataset S3) associated with a positive effect size value (thus cavity) contained a 3-bp deletion as compared with the opposed unitig (n126874), which reduced the number of proline repeats from 4 to 3 in the C-terminal region of Pks13 (Fig. 5B). Structural alignment (Fig. S3) revealed that this proline repeat is unique to *M. avium* homologues and is located between the C-terminal acyl carrier protein (C-ACP) domain and the thioesterase (TE) domain of *M. tuberculosis* homologue (48). Unitigs with equally low *P* values were clustered only over the Pks13 coding region (Fig. 5A). These findings suggest the possibilities of the association of linkage disequilibrium with disease type. Therefore, we inferred that the Pks13 variant could also contribute, at least in part, to the bacterial phenotype, leading to cavity formation.

**FIG 5.**
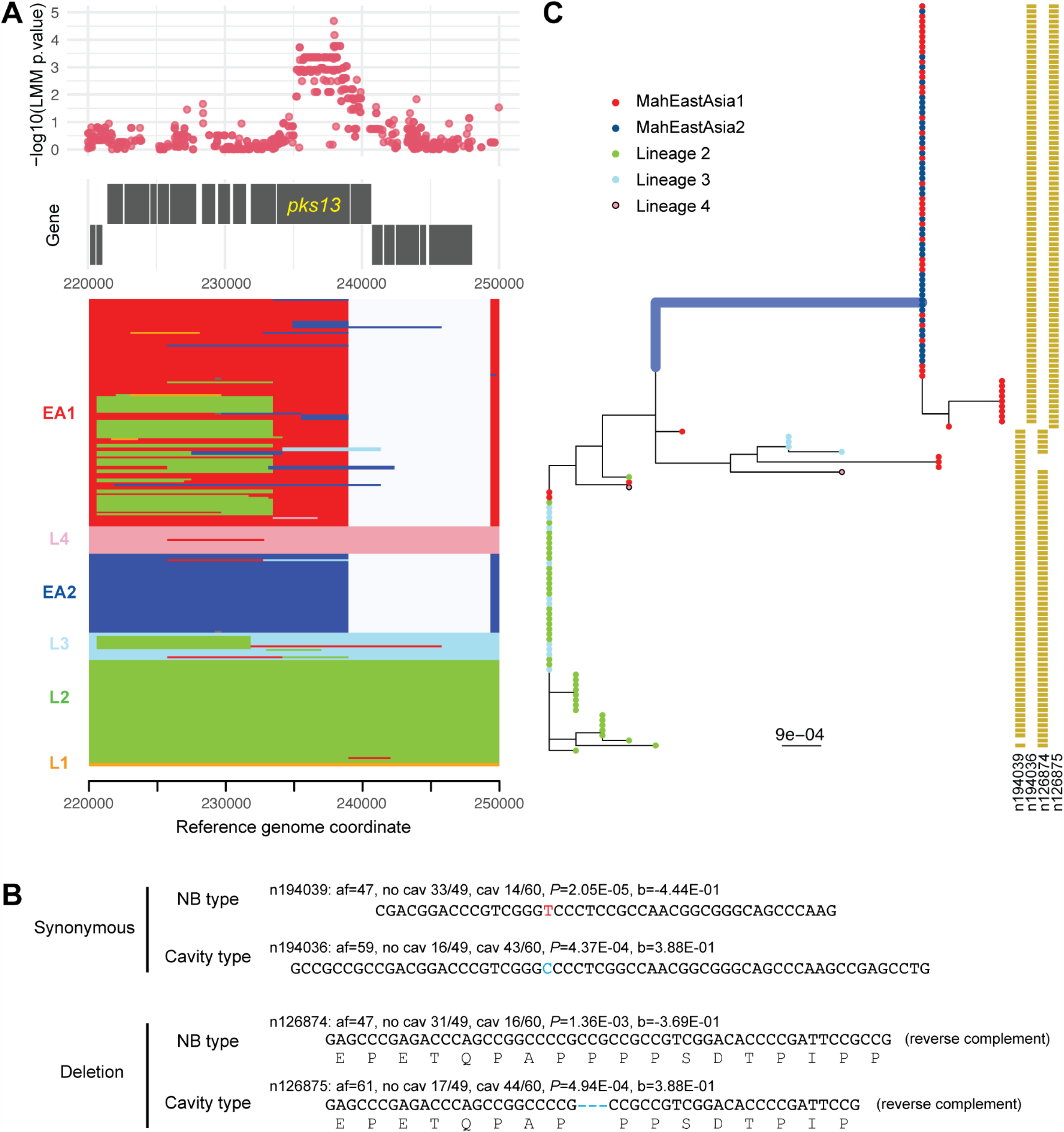
Allelic variation in the coding region of Pks13. (**A**) Upper panel: locus effect *P* values around coordinate 235,000. Lower panel: inter lineage recombinations detected within reference genome coordinate 220,000 to 250,000. The color indicates donor lineage except for grey. Grey indicates ancestral recombination between EA1 and EA2. (**B**) Unitig pairs containing synonymous substitution or deletion. (**C**) Phylogenetic tree of Pks13 variants. The alignment was generated using gene products from recombination free MAH_0232 equivalent locus of 148 global strains. The tree was constructed based on the LG model using FastTree v 2.1.0 (70). The branch acquired cavity-associated motifs are highlighted by a thick line whose length is equivalent to 10 amino acids change. Tip color indicates the lineage of the strain. The side columns indicate the presence of the four unitigs in the respective coding region.

The unitig causing proline deletion in Pks13 (n126874) was found in 41 of 43 EA2 members and 50 of 124 EA1 members (Dataset S2). The recombination tracts in the EA1 genomes cover a large portion of the coding region, wherever detected (Fig. 5A). Inter-lineage recombination was rare in the EA2 genomes, whereas ancestral recombination shared between EA1 and EA2 was detected downstream of the significant unitigs (Fig. 5A). Phylogenic inference for *M. avium* Pks13 variants, based on recombination-free sequences, showed advanced differentiation of EA2 variants from the lineage 2 variants (Fig. 5C). Cavity-associated two unitigs (n194036/n126875) were present in the coding regions of two or all three variants from EA1/EA2 members connected to the highlighted long branch in Fig. 5C. These findings indicate that the cavity-associated unitigs (n194036/n126875) and the resulting product variants might have originated from the EA2 lineage and spread to a subset of EA1 members through recombination.

### ‘No cavity’-associated locus missing in the reference chromosome

The DBGWAS analysis revealed a significant unitig present only in the no cavity group (n2184, af = 8, no cavity 8/49; cavity 0/60, *P* = 9.62e-06, effect size = -8.19e-01, Comp_0 in Dataset S3). The sequence motif, a part of the hypothetical protein gene, was located on a contig of strain MK20-1 draft genomes (accession no. NZ_BDNW01000006) (18) but was missing in the strain TH135 chromosome. These data imply a plasmid or genomic island in the *M. avium* population, which could have a negative impact on cavity formation.

### Distribution of cavity risk variants in the global population

The distribution of cavity risk variants in the three chromosomal loci is summarized in Fig.1B and Dataset S2. After filtering the strains with minor or ambiguous risk variants detected in Osaka strains, 33 (80.4%) out of 41 EA2 members carried the complete set of cavity risk variants. Twenty (18.8%) out of 106 EA1 members carried the complete set. Thirty-two (96.9%) out of 33 filtered lineage 2 members carried only NB (no cavity) -associated variants at all three loci. None of the lineage 2 members carried the complete set of cavity risk variants.

## DISCUSSION

In this study, using the bacterial GWAS approach, we have identified three cavity-associated loci (*ftsHp, pks13*, and CYP139) whose polymorphisms have not been mentioned in previous comparative and functional genomics studies of *M. avium* (18, 30, 31). The frequent occurrence of these three sets of cavity risk variants in East Asian lineage genomes is consistent with the high prevalence of NTM lung disease in Asia (8), suggesting that the bacterial genetic variation and the resulting phenotypic variation among *M. avium* clones influence the epidemiology of NTM lung disease. Furthermore, retrospective studies on chronic *Mycobacterium* lung infection have shown the emergence of various antimicrobial resistance mutations in *M. tuberculosis* and *M. abscessus* clones during therapy (49, 50). However, except for the 23S rRNA gene, none of the three loci-equivalent regions was identified as hot spots of resistance mutations in previous retrospective studies (49, 50). Therefore, the variants in the three loci identified in this study are likely unrelated to bacterial adaptive mutations that occurred during therapy.

FtsH is a membrane-anchored zinc metalloprotease that presumably acts on membrane proteins (51). *ftsH* expression is induced under oxidative stress in *M. tuberculosis*, and its control expression is important for cell growth in macrophages (52). However, neither *ftsH* nor *fol* genes are essential for the growth of *M. avium* strain 11 or MAC 109 (30, 31). Therefore, we speculate that the 39 bp repeat variant in the *ftsH* promoter region could influence not only the activity of the folate synthesis pathway but also the stress resistance through the high expression of FtsH protease in the host.

In general, *Mycobacterium* possesses a large variety of cytochrome P450 (41). The genome of *M. avium* strain 104 carries 48 cytochrome P450 loci, covering 28 distinct P450 families. The MAH_2197 locus (equivalent to MAV_3106 in strain 104 and DF55_111000 in strain MAC109) is the only locus encoding the CYP139 family protein in the *M. avium* genome. The CYP139 family, one of the P450 families with the top eight lowest evolutionary rates (41), was present across all *Mycobacterium* species analyzed in the previous study. Despite being highly conservative, CYP139 has been suggested to be nonessential for *M. avium* growth through *in vitro* (30) and *in vivo* (31) studies using MAH strain alone, and with mice, respectively. Although the exact substrate of the CYP139 family protein is unknown, CYP139 is postulated to be involved in producing a secondary metabolite since the coding sequence of the CYP139 family protein is located within a biosynthetic gene cluster (53). In the *M. avium* genome, the CYP139 gene is next to the type III polyketide synthase gene. Therefore, more attention should be paid to CYP139 regarding its substrate, product, and influence on the host.

Furthermore, the GWAS results highlighted polymorphisms in Pks13. However, the effects and mechanisms of the synonymous substitution polymorphism in pks13 on the bacterial phenotype are unclear. The cluster of unitigs with equally low *P* values implies that a set of amino acids on the uniquely diversified Pks13 variants of EA1/EA2 members also contributes to the risk of cavity formation. Pks13 is responsible for the condensation process that connects the α-alkyl chain and the meromycolate chain of mycolic acid in the mycolic acid assembly and mycolate transfer to trehalose in TMM biosynthesis (42, 46, 47). Psk13 is essential in *M. smegmatis* (42), *M. tuberculosis* (54), and *M. avium* strain MAC 109 (lineage 3) (30), but not in *Corynebacterium* (42), and interestingly *M. avium* strain 11 (lineage 3) (31). Therefore, there is an obvious variation in the intrinsic cell wall integrity between the *M. avium* strains. Furthermore, the structure of the meromycolate chain differs among bacterial species (55). It can be inferred that the variation in the Pks13 sequence might reflect the difference in the cell envelope structure.

Together, this study identified three loci involved in lipid synthesis (Pks13 and CYP139) or membrane protein maintenance (FtsH), suggesting that natural variation in biosynthesis or maintenance of *M. avium* membrane components influences the host disease type of MAC lung disease. The direct cause of the cavity is thought to be the host matrix metalloprotease acting on granuloma (56). Granuloma is induced by mycobacterial trehalose 6, 6’-dimycolate (TDM) (57), which is exported outside mycobacterial cells through MmpL3 protein embedded in the plasma membrane (58). Therefore, the implication of the Pks13 variant and membrane-anchored protease FtsH in cavitary disease is congruent with the current model for cavity formation in mycobacterial lung disease.

The major limitations of this study are the small sample size and the inability to arrange bacterial strains before antimicrobial therapy, especially for the cavity group. Thus, the false positives and variants associated with antimicrobial tolerance may underlie the *P* value peaks. In addition, we could not explain the biological significance of the significant unitigs with synonymous substitutions in *psk13* (MAH_0232) and dihydrodipicolinate reductase gene (MAH_3077). Nevertheless, we emphasize that this GWAS focused on East Asian lineages could rank the polymorphic sites in the *M. avium* genome based on the strength of association with cavity formation. The identified variants at high ranks are not just bacterial genetic markers for prognosis prediction of MAC lung disease, but one can also be the targets for discovering new drugs (43–45) for NTM lung disease. This study provides mechanistic insights into *M. avium* virulence, and therefore, could serve as the foundation for future personalized medicine. However, future GWAS on different *M. avium* subpopulations are necessary to validate the findings of this study, including the implication of non-core genes.

## MATERIALS AND METHODS

### Bacterial strains

All the 109 MAH strains used in GWAS were isolated from sputum specimens of immunocompetent adult (> 42 years old) patients visiting the National Hospital Organization Osaka Toneyama Medical Center (N 34.793281 E 135.453215). The isolates were cultured on Kyokuto 2% Ogawa medium (Kyokuto Pharmaceutical Industrial Co., Ltd., Tokyo, Japan) and BD Difco™ Mycobacteria 7H11 agar supplemented with BD BBL™ Middlebrook OADC enrichment (Becton, Dickinson and Company, NJ, USA). Metadata linked to 109 strains are summarized in Dataset S1.

### Metadata linked to bacterial strains

Among the 109 patients with lung disease, 108 met the diagnostic criteria of MAC lung disease recommended by the American Thoracic Society/European Respiratory Society/European Society of Clinical Microbiology and Infectious Diseases/Infection Disease Society of America (ATS/ERS/ESCMID/IDSA) (28), whereas the remaining one patient did not qualify, as the culturable *M. avium* could only be detected once from the patient’s sputum. Disease type, duration of disease, and history of antimicrobial therapy (the treatment regimen comprising two or more antimicrobial drugs) were collected from the medical records of the patients. The chest computed tomography (CT) findings were used to decipher the disease types (FC, NB, NB with cavity, and unclassifiable disease type without cavity). The patients who have remained ‘NB alone’ at least for five years without or with antimicrobial therapy were included in the NB category. FC disease was defined as the presence of cavity and fibrosis in the upper lobes of the lungs as the main finding, while NB disease was characterized by the presence of multicentric nodules and bronchiectasis. Two pulmonologists evaluated the radiographic findings and serial changes independently, and the final diagnosis was determined by consensus.

### Antimicrobial sensitivity tests

The MIC of sulfamethoxazole/trimethoprim mix (19:1 ratio) for MAH strains were determined using Kyokuto Opt Panel MP (Kyokuto Pharmaceutical Industrial Co., Ltd.), designed to perform the broth microdilution following the guidelines in the Performance Standards for Susceptibility Testing of Mycobacteria, Nocardia spp., and Other Aerobic Actinomycetes, 3^rd^ edition (document M24) of the Clinical and Laboratory Standards Institute (CLSI). The sulfamethoxazole/trimethoprim concentrations (µg/ml) tested were 4.8/0.25, 9.5/0.5, 19/1, 38/2, 76/4, and 152/8. The test strains were precultured in Mycobroth (Kyokuto Pharmaceutical Industrial Co., Ltd.) for a week, and the turbidity of the culture was adjusted to an absorbance of 0.08–0.10, using a spectrophotometer/fluorometer (DS-11FX+, DeNovix, USA). The bacterial suspension was diluted 100-fold with Mueller-Hinton broth (Kyokuto Pharmaceutical Industrial Co., Ltd.) supplemented with 5% (v/v) Middlebrook OADC enrichment, inoculated into Kyokuto Opt Panel MP, and cultured at 37°C for seven days. The antimicrobial potency was based on an 80% growth inhibition.

### Preparation of genomic DNA

Strains were propagated on 7H11 agar medium supplemented with Middlebrook OADC enrichment and incubated for up to three weeks at 37°C. DNA samples for sequencing were prepared from the pooled colonies grown on the agar plates using either the traditional method that combined enzymatic lysis and bead-beating (59) (five samples denoted as ‘mechanical_ProK’ in DNA_extraction column in Dataset S1) or more quick mechanical lysis only protocol shown in Text S1 (100 samples denoted as ‘mechanical_only’’ in the DNA_extraction column in Dataset S1).

### Sequencing and genome assembly

Of the 109 strains, the genome sequences of four strains, OCU901s_S2_s2, OCU404, OCU464, and OCU466, were released without linked metadata in a previous study (19). Therefore, in this study, paired-end reads of 105 strains were obtained using the Illumina platform. Library construction and sequencing were performed in Macrogen (Macrogen Japan Corp, Tokyo, Japan; HiSeq 2500), BGI (BGI JAPAN, Kobe, Japan; HiSeq 2500), or University of Minnesota Genomics Center (Minneapolis, MN, USA; NovaSeq 6000). The Illumina TruSeq DNA PCR-Free Library Kit (Illumina, Inc., CA, USA) was used to prepare DNA libraries. More than 1 Giga base-pair information was obtained per sample. Paired-end reads were trimmed using fastp software v0.19.5 (60) using the command ‘fastp -i ../$name1 - I ../$name2 -3 -M 30 -c -o ./$prefix_R1.fastq.gz -O ./$prefix_R2.fastq.gz - h ./$Prefix_report.html -j ./$prefix_report.json -q 30 -n 10 -t 1 -T 1 -l 20 -w 16’, then the trimmed reads were assembled using Unicycler v0.4.8 (61) with the command ‘unicycler -1 ../../fastp/$name1 -2 ../../fastp/$name2 -o unicycler_out -t 16 --keep 0’. Unicycler assemblies of 105 strains and previously published assemblies of four strains (OCU464, OCU466, OCU404, OCU901s_S2_2s) were used as inputs for GWAS. Genome assemblies are available from figshare (62).

### Inference for genetic population structure

A population genome dataset of 255 MAH genomes was created by combining 150 genomes available from the RefSeq database at Oct 29 2020 and newly sequenced 105 genomes to infer the lineage to which each strain belongs. Core genomic SNPs were detected by Parsnp and HarvestTools (63) using the chromosome sequence of strain TH135 (accession no. AP012555.1) as the reference genome. Command used was ‘parsnp - g ../Mah_TH135.gbk -d ./fasta_MAH_255_minus_TH135 -p 8 -c’. Polymorphic sites containing N were excluded. To infer chromosomal recombination in fastGEAR, filtered SNPs (PASS call in Parsnp output) were combined with intervening reference genome sequences per genome, then a multi-FASTA file of 4,949,973 bp long 255 genome sequences with 43,526 polymorphic sites was created. Assuming that MAH is a recombining species, partitioning the global population into sequence clusters (SCs) was inferred using the BAPS3 program implemented in fastGEAR (37). SC is a subpopulation treated as a random mating unit, inferred without using SNP position information. Lineage in the global population was defined based on the proportion of fragment length sharing the ancestry (PSA) between the two genomes. Based on the recombination/lineage inference results of fastGEAR, another round of PSA calculations was performed using fastGEAR output files and an R script. Scripts and data files used for visualization are available from figshare (64).

### GWAS

We used the DBGWAS pipeline (38) to conduct a reference-free GWAS based on a linear mixed model. DBGWAS relies on genome assembly & analysis tool box (GATB) (65) for k-mer detection and their compaction into unitigs and bugwas (66) for detecting significant associations between unitigs and phenotypes. As an objective variable of the phenotype, the cavity was set to 1 (n = 60), while no cavity was set to 0 (n = 49). A phylogenetic tree based on core genomic SNPs on 255 genomes produced by Parsnp was used for lineage effect analysis. We focused on unitigs with a *P* value < 10^−4^ and their neighboring unitigs in the network graph outputs from DBGWAS. DBGWAS was conducted using a singularity image file with the command ‘singularity run ../../../software/dbgwas-0.5.4.simg -strains ./strains_strict_ME_added2.txt - newick ./parsnp.newick -maf 0.5 -nc-db ./MAH_gene_db.fas -pt-db ./uniprot_sprot_bacteria_for_DBGWAS.fasta’. Unitig location on the reference chromosome was identified by conducting the blastn search (67) using all unitig sequences as queries and the chromosome sequence of strain TH135 as the subject.

### Phylogenetic analysis

A phylogenetic tree of 255 strains (Fig. 1) was constructed based on the p-distance matrix generated from the SNP alignment produced by Parsnp (63). For the phylogeny construction of Pks13 variants, the orthologs of MAH_0232 products in 255 genomes were identified using the tblastn program (67) using the MAH_0232 product as a query, and the hit coding sequences were retrieved and aligned using MAFFT v7.407 (68). Multi-FASTA entries with recombination tracts were removed using RPF4 (69). A phylogenetic tree of Pks13 was constructed using the approximately-maximum-likelihood method on FastTree v. 2.1.0 (70) based on the alignment of the translated sequences. Tree and genotype data were visualized using the ggtree package of R (71). Scripts used, and tree with tip labels are available from figshare (64, 72)

## ETHICAL STATEMENTS

The National Hospital Organization Osaka Toneyama Medical Center review board approved the study protocol (approval number TNH-2018035-4). The Kobe Institute of Health review board approved the study protocol (approval number Sen2-3).

## DATA AVAILABILITY

Raw sequence reads generated in this study are deposited in DDBJ Sequence Read Archive under accession number DRA011807. Genome assemblies used are available from figshare DOI: https://doi.org/10.6084/m9.figshare.14788857. R scripts used for data analysis and visualization are available from figshare DOI: https://doi.org/10.6084/m9.figshare.14788881.

## ACKNOWLEDGEMENTS

We thank Shusei Sato (Graduate School of Life Sciences, Tohoku University) and Masaru Bamba (Graduate School of Life Sciences, Tohoku University) for discussions on GWAS results and Yukari Sato (Graduate School of Life Sciences, Tohoku University) for instructions on the PyMol. We thank Manabu Ato (National Institute of Infectious Disease Japan) and Ho Namkoong (Keio University School of Medicine) for supporting the project administration of the Japan Agency for Medical Research and Development (AMED) grant. We also thank Editage [http://www.editage.com] for editing and reviewing this manuscript for English language. The computation was supported by the Institute of Medical Science, the University of Tokyo, Human Genome Center supercomputer system SHIROKANE.

This research was supported by AMED grant numbers JP21fk0108621 (H.Y., K.A.) and JP20fk0108129 (F.M., H.K., Y.N., T.I.), JP19fk0108043 (F.M., M.M., Y.N., T.I.), JP18fk0108043 (S.K., F.M.), and in part by Japan Society for the Promotion of Science (JSPS) KAKENHI grant numbers 18K06357 (H.Y.), JP21H02093 (H.Y.), JP19K08629 (Y.N.), and JP18K10041 (T.I.). The funders had no role in the study design, data collection and interpretation, or the decision to submit the work for publication. Author contributions are as follows: H.Y., S.K., and T.I. conceptualized and designed the research; S.K. and M.M. collected clinical metadata; S.K., M.M., and H.K. provided bacterial strains; F.M. and H.K. administrated the project; Y.N., K.A., A.O. and T.I. isolated strains and prepared genomic DNA; Y.N. performed MIC experiments; H.Y. conducted formal analyses, data curation and visualization; H.Y., Y.N., S.K. drafted the manuscript; H.Y., Y.N., F.M., and T.I. reviewed and edited the manuscript.

## Supplemental Materials

**Fig S1** (FIG S1.pdf)

Recent recombination and the PSA matrix. (A) Complete linkage clustering of 255 strains and visualization of recent recombination. Distance is defined by 1-PSA. (B) The PSA matrix.

**Fig S2** (FIG S2.pdf)

Alignment of *ftsH* promoter region. The intergenic region between MAH_0464 and MAH_0465 (*ftsH*) is shown. Polymorphic sites were also highlighted, and the 39 bp repeats are underlined.

**Fig S3** (FIG S3.pdf)

Alignment of Pks13 homologs. Alignment was generated based on the secondary structure using the PROMAL3D web server (75). The TH135 homolog is a cavity-risk variant. The domains are indicated above the alignment.

**Dataset S1** (Dataset S1.xls)

*M. avium* Osaka strains and the linked metadata.

**Dataset S2** (Dataset S2.xls)

Distribution of cavity risk variants in global *M. avium* population.

**Dataset S3** (Dataset S3.xls)

Nucleotide sequences of unitig-network graph components. The information was selected from the original DBGWAS output. The output of DBGWAS, including graphical summary, is available from fighshare (64) under ‘DBGWAS/visualization/data/output/visualisation’.

**Text S1** (Text S1.docx)

Genomic DNA extraction from *M. avium*: mechanical lysis protocol.

